# Talent Identification at the limits of Peer Review: an analysis of the EMBO Postdoctoral Fellowships Selection Process

**DOI:** 10.1101/481655

**Authors:** Bernd Klaus, David del Álamo

## Abstract

Scientific peer review is still the most common system for fund allocation despite having been shown in multiple instances to lack accuracy in identifying the most meritorious applications among high quality ones. This study evaluates two aspects of the selection process of the top-ranked applicants to the EMBO Long-Term Fellowship program in 2007. First, the accuracy of the system is evaluated by comparing the level of career progression of the candidates in 2017 with the original award decisions made in 2007. The second aspect, explores the relationship of career progression with indicators derived from the information available to evaluators at the time of application. The results obtained suggest that the peer review system is not substantially better than random selection in identifying the best candidates once an initial pre-selection of the most promising ones is performed. Not only that, the analysis of the indicators studied, some of which have not been analyzed in detail in the past, suggests that among other potential sources of uncertainty, the information available at the time of application is not sufficiently predictive of career progression. As previously described, however, we find clear differences in career progression between men and women. We propose a new mixed model of fellowship evaluation in which peer review is used to select high quality applications, and random allocation of funds is subsequently used to award fellowships among these top ranked candidates.

## Introduction

Peer review evaluation is widely established in academic scientific environments and, with variations, the process is straightforward: experts in a given area assess the professional performance or quality of individuals, project proposals or scientific production in their own field of competence. The system is based on the assumption that active researchers are the best judges of the results and performance of other scientists in their field of competence [1, 2]. Indeed, numerous analyses in the past decades have suggested that peer review selection is able to reasonably discriminate between high- and low-quality proposals [3–5].

However, evidence arising from the evaluation of the outcome of research grants selection processes and, less frequently, from the selection of fellowship awardees, suggests that the peer-review selection system is less than optimal in identifying high potential in research projects or scientists, respectively, if the decision has to be made within a group of pre-selected, high-quality applications [1, 4, 6–8]. Peer review procedures “usually succeed in identifying flawed or conceptually weak proposals. However, decisions concerning ranking and funding level of more competitive proposals can be markedly subjective and opaque” [9; p. 2]. Evidence suggests that peer review fails to perform in finely discriminating those that have a true potential from those of a lesser value among high-quality applications [10–13].

We analyze here the peer review selection process of the EMBO Long Term Fellowship (ELTF) programme. Active for more than 50 years, this prestigious international programme funds postdoctoral researchers in the area of molecular biology (and more recently in broader areas of biology) for a period of up to two years (http://embo.org/funding-awards/fellowships/long-term-fellowships, last access November 2018). Selection of candidates takes place in two peer review steps: a pre-selection step, in which ~1/4 of the candidates are pre-selected and a second selection round in which about ~1/2 of these preselected candidates are awarded a fellowship. A committee formed by active scientists makes the decisions in both steps.

As observed in other funding schemes [see for instance 14], ELTF success rates have dropped significantly over the last 40 years, from a success rate of ~40% in the late 1970s to an average of 14% in the period 2013–2017 (EMBO Facts and Figures 2017, http://www.embo.org/documents/news/facts_figures/EMBO_facts_figures_2017.pdf, last accessed October 2018). Similar trends have been observed in other funding programs, and are usually caused by an increase in the number of applications combined in some cases with reduced availability of funds [see for instance 2]. Considering these success rates, acceptance and rejection decisions are being made inevitably within a pool difficult to rank where differences between candidates are minimal. In the words of S. Vazire, “the fact is that separating shoddy work from solid work is much more straightforward than distinguishing the top 5% of solid work from the next 5%, which often makes the difference between favourable and unfavourable decisions” [15; p. 7].

A previous analysis of the ELTF programme based on bibliometric indicators (number of articles and citations received) showed that, considering all applicants of a given year, the subsequent productivity of the awarded candidates is statistically higher than that of the rejected ones [4]. This is in line with studies of other selection processes suggesting that peer review is able to gross discriminate among an entire range of proposals [9, 12].

We focused, however, in this study on the candidates that were pre-selected after the first step of evaluation, those who in light of the evidence discussed above should be difficult to rank. In particular, we focused on decisions made among already pre-selected, high quality applications to the ELTF programme in 2007 and analyzed whether these decisions resulted in awards to the most promising candidates as judged by their performance ten years later, in 2017. We also analyzed the relationship between future performance and two sets of indicators based on the information provided by the candidates in 2007 (see the **Methods** section for details): traditional productivity or impact-based scientific indicators, such as publications, citations and Journal Impact factor, and “social indicators”. As explained above, fine discrimination among high quality applicants is not straightforward and may lead to decision bias [16]; “when there is no objective basis for choosing one qualified candidate over others, people naturally fall back on subjective preferences. A selection committee might consciously or unconsciously favour certain research topics, groups of people or even individuals” [15; p. 7]. Social indicators analyzed include the prestige of former and future supervisors, the prestige of former and future scientific institutions where the candidate has worked or will work and the effect of the country of nationality of the candidate as well as the countries where they have worked and plan to work in the future.

In order to compare candidates with one another based on subsequent performance, a measure of performance in 2017 is required. In the absence of a universally accepted measure of scientific merit [4, 17], we combined two aspects of the scientific activity to create a single measure of performance (Pf) to rank the candidates in this study. One aspect is scientific productivity (number of publications and citations received), which has been traditionally used as a measure of scientific performance [see for instance 4, 18, 19], and the other aspect takes into consideration career progression according to the *standard* scientific academic career: “The ideal scenario for a postdoctoral researcher’s experience might be as follows. A self-selected, motivated, recent doctoral degree recipient receives additional scientific training (to augment previous training during graduate school) for a limited time before transitioning to a full-time research position, often as a tenure-track faculty member” [20; p. 17]. In this regard, the time the candidate has spent as an independent researcher is also taken into consideration. Pf is therefore a proxy for scientific career progression calculated as a combination of articles published, citations and time the candidate spent as a group leader if at all, all measured ten years after application to the ELTF programme (see **Methods** for details on the precise calculation of Pf).

Our results show that, in line with previous reports, peer review failed to discriminate the candidates that later on showed significant career progression from those who did not. Mathematically, peer review selection did not demonstrate any advantage over random selection.

Moreover, none of the indicators, scientific or social, extracted from the data available in 2007 correlate significantly with career progression (see **Results**). We found, however, a clear correlation of gender with career progression: male applicants tend to progress significantly more than their female counterparts in their academic careers, as previously observed in multiple studies.

## Methods

### Data description

1288 early career researchers (45.2% females) applied to the LTF programme in 2007 and 533 were pre-selected for the second step of selection. 212 candidates were awarded the fellowship. The initial working group is composed by the 212 candidates awarded (38.7% females) and the next 212 candidates (46.7% females) in the ranking that were not awarded the fellowship as the control group. From these we excluded 35 candidates that could not be unequivocally identified in 2017, and 62 candidates (51.6% females) who abandoned active research and could therefore not be ranked according to Pf. The working group analyzed in this report is composed by 327 individuals (40.4% females), candidates to an ELTF in 2007 and still active researchers in 2017. 172 of them (52.5%) were awarded an ELTF.

### Performance (Pf)

The group of 327 researchers was ranked according the value of Pf. Pf is composed of the following factors:

- Time in years that the researcher has spent as group leader (0–10). Differences in months are not taken into consideration, as precise information was not available in most cases. Information was manually collected through internet searches between the months of March and April of 2017.
- Number of research articles published as corresponding author by the researcher as group leader. Only primary research articles published in English in international peer-reviewed journals listed in the PubMed database were taken into consideration (PubMed, National Library of Medicine (US). https://www.ncbi.nlm.nih.gov/pubmed/, data collected between March and June 2017).
- Average number of citations per year of these articles. This information was obtained from the Core Collection Database of Thompson Reuters’ Web of Science (Web of science™, Thompson Reuters http://apps.webofknowledge.com/WOS_GeneralSearch_input.do?product=WOS&search_mode=GeneralSearch&SID=N2Tma68dipAlrbGYZuT&preferencesSaved=, data collected between March and June 2017).

Data in each category was normalized using z-scores and Pf was calculated as the average of the z-score of each researcher in each category. Please note that the actual value of Pf (ranging from -0.70 to 4.22) has no intrinsic meaning and is used in this context to compare the candidates with one another in terms of career progression.

Candidates were binned into four classes (Pf classes) by k-medoids clustering (a variant of k-means, see Supplementary Figure 1).

Pf classes are composed as follows:

- Group Leader–High (GLHi): 66 individuals, group leaders for 6 years on average. They published on average 6.1 articles as corresponding authors in this period and these articles were cited 5 times per year on average.
- Group Leader-Medium (GLMe): 64 individuals, group leaders for 4.7 years on average. They published on average 2.1 articles as corresponding authors in this period and these articles were cited 2 times per year on average.
- Group Leader-Low (GLLo): 74 individuals, group leaders for 2.6 years on average. They published on average 0.2 articles as corresponding authors in this period and these articles were cited 0.7 times per year on average.
- Non-independent researchers (NI): 123 individuals not yet group leaders as the time of analysis, but still active as researchers.

### Scientific indicators

Four bibliometric independent variables measured at the time of application were analyzed for their effects on Pf class distribution of candidates: total number of articles published by the candidate, total number of articles as first (or co-first) author (referred to as “first author” for simplicity), average Journal Impact Factor (JIF) of the journals where the first author articles were published and total sum of the JIF of these journals. Number of articles published was provided at the time of application and information was complemented using the PubMed database (see above).

*JIFs* were obtained through Thompson Reuters’ Journal Citation Reports (InCites™ Journal Citation Reports^®^ 2015, Thompson Reuters, https://jcr.incites.thomsonreuters.com/JCRJournalHomeAction.action, For simplicity, the latest available JIFs at the time of analysis (2015) were used; a comparison between the JIFs of 65 of the most commonly used journals in molecular biology and related areas in 2006 and 2015 reveals substantial changes in the actual values, but very minor changes in the relationships between journals: ranking of journals according to their JIF is very similar in 2015 compared to 2006 (data not shown). Note that JIF is nowadays widely discredited as a measurement of performance [21–23], but its use as a proxy for individual performance was not uncommon in 2007. Even today, despite heavy criticism, its presence pervades peer-review evaluation systems [see for instance 24].

### Social indicators

Three different categories of social variables were analyzed:

- *Supervisor network size*. Network sizes of the candidate’s PhD supervisor and the proposed host supervisor (the researcher with whom they propose to develop the project described in the application) were estimated using the number of unique co-authors of the supervisors in articles published in the two years prior to the first deadline for application in 2007 (15/02/2005 to 15/02/2007) as a proxy. Evidence suggests that network size can be reasonably estimated based on the number of coauthors [25, 26]. Number of co-authors was manually calculated taken into consideration all types of publications listed in the PubMed database in the period listed above.
- *Institutional prestige*. Although institutional prestige is obviously a subjective quality, a proxy for prestige that is commonly used in the literature is institutional rankings [18, 27–29]. In this study, ranking of the institution where the candidate obtained her or his PhD and that of the host institute where they proposed to work was based on data from the Mapping Scientific Excellence online application (Mapping Scientific Excellence, www.excellencemapping.net, last accessed June 2017). Institutions were ranked according to the *best journal rate* in *Biochemistry, Genetics and Molecular Biology* subject area (no covariates) based on the publication period 2009–2013 [see 30, for further details].
- Scientific ranking of the country of nationality of the applicant, country where the candidate obtained his or her PhD and country where the proposed host institution is located. Country ranking was based on the SCImago public database (SCImago Journal & Country Rank, retrieved July 21, 2015, http://www.scimagojr.com, last accessed June 2017). Countries were ranked according to the productivity indicator *h-index* [31] in the subject area *Biochemistry, Genetics and Molecular biology*.

### Statistical analysis

Data collection and preliminary statistical analysis (Pf calculation and ranking of candidates) was performed using Microsoft^®^ Excel^®^ for Mac 2011.

Further statistical analysis was performed using R, an open source programming language for statistical computing [32; see also https://www.R-project.org/, R Foundation for Statistical Computing, Vienna, Austria)]. Details are described in the **Results** section where applicable. An Rmarkdown document (html) containing the complete data analysis details is available at https://git.embl.de/klaus/embo_ltf_analysis and as supplementary material.

## Results

Previous studies suggest that peer review evaluation of already pre-selected, high quality candidates would not be able to detect the most promising vs. other highly qualified individuals. In our case, a selection process with high predictive value should be able to award the fellowship to a population highly enriched in individuals from the GLHi class, and deny the award to individuals mainly grouped in the NI class as defined in the **Methods** section. However, data represented as a dot plot in Figure 1A showed a rather similar distribution in Pf classes of awarded vs. rejected candidates. This result suggests that the peer review process failed at fine ranking of applicants that were already pre-selected.

**Figure 1.**
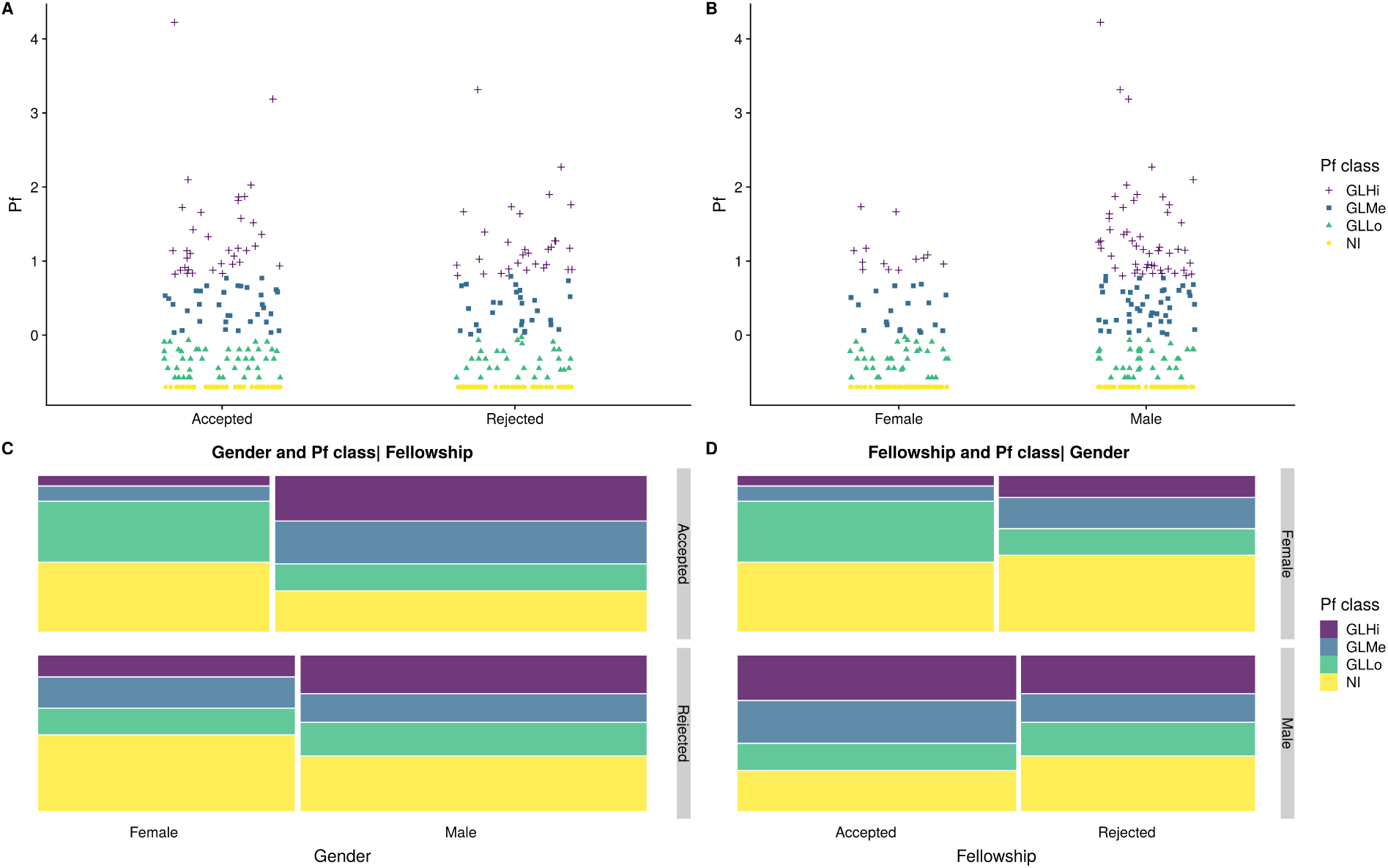
Relationships between Gender, Performance and acceptance into the fellowship. (A) Performance factor for accepted and rejected candidates respectively. (B) Performance factor for women and men. (C) Relationship between gender and Pf class given the fellowship status. (D) Relationship between fellowship status and Pf class given gender.

A rather different picture emerges when the applicants are distributed into the Pf classes according to their gender (Figure 1B). Results showed a clear difference in female vs. male applicants: females tended to populate the NI class in a bigger proportion than males and reciprocally, males are enriched in the GLHi class compared to females. This result is expected in light of the very well documented gender inequality that still persists in scientific careers (see the **Discussion** section). Out of the 66 individuals contained in the GLHi group, only 13 were female researchers. Correspondingly, although the numbers were similar for males and females in the NI group (60 vs. 63), these represented a much smaller proportion of the total number of males (195 vs. 132). In a log-linear regression model of homogeneous association using the Pf class, gender and fellowship status (awarded or rejected), gender and class were not independent given fellowship status (For the terminology, see [33], Tables / Figure 8.1 / 8.2 and Eq. 8.11 in section 8.2). The differences in Pf class distribution of applicants according to gender given the fellowship status was borderline statistically significant as evidenced by a log-linear regression model (Likelihood-Ratio (LR)-Test, P=0.1072, Figure 1C - Note that we use a Quasi-Poisson model by default in the LR tests in order to account for over-dispersion and obtain reliable p-values). A simplified analysis taking into consideration only the most extreme classes, GLHi vs. NI, also shows that males populate the GLHi class in a much higher proportion than females and that this difference is highly significant (LR-Test, P=0.00741). In contrast, using the log-linear regression model to control for the influence of gender, no statistically significant difference was found in the distribution into Pf classes of accepted vs. rejected candidates (LR-Test, P=0.87020). This is graphically represented by the mosaic plot in Figure 1D. As a control, we did not find any statistically significant association of gender with acceptance or rejection of the fellowship given the class label (LR-test, P=0.72605, Supplementary Figure 2), indicative that the differences between males and females arise later on in their careers. Interestingly though, we found a specific effect for females: there was a particularly large proportion of GLLo female group-leaders (see Figure 1C and Supplementary Figure 2), which may reflect a delay in their career progression with respect to their male counterparts.

In conclusion, our data indicates that the peer review selection system of ELTF is unable to identify the most promising candidates, as there is no statistically significant difference in career progression between candidates awarded and candidates rejected. There is however a profound difference between female and male candidates. Gender, as described in multiple previous studies, is still sadly a good predictor of career progression.

### The effect of scientific indicators on career progression

Scientific indicators considered in this study are bibliometric factors traditionally used in scientific research assessment, measured at the time of application in 2007. As explained in the **Methods** section, the following indicators were tested: total number of publications; number of publications as first author; average *JIF* of the journals where first author publications were published; and total sum of the *JIF* of the journals where first author publications were published.

Figure 2 shows the dot blot representation of the distribution of applicants in Pf classes according to the total number of publications (Figure 2A) or the number of first author publications (Figure 2B). While no pattern is obvious in the representation there seems to be an association between a higher number of total publications and a higher Pf class. This association is more obvious when a mosaic plot is used instead (Figure 3A and 3B).

**Figure 2.**
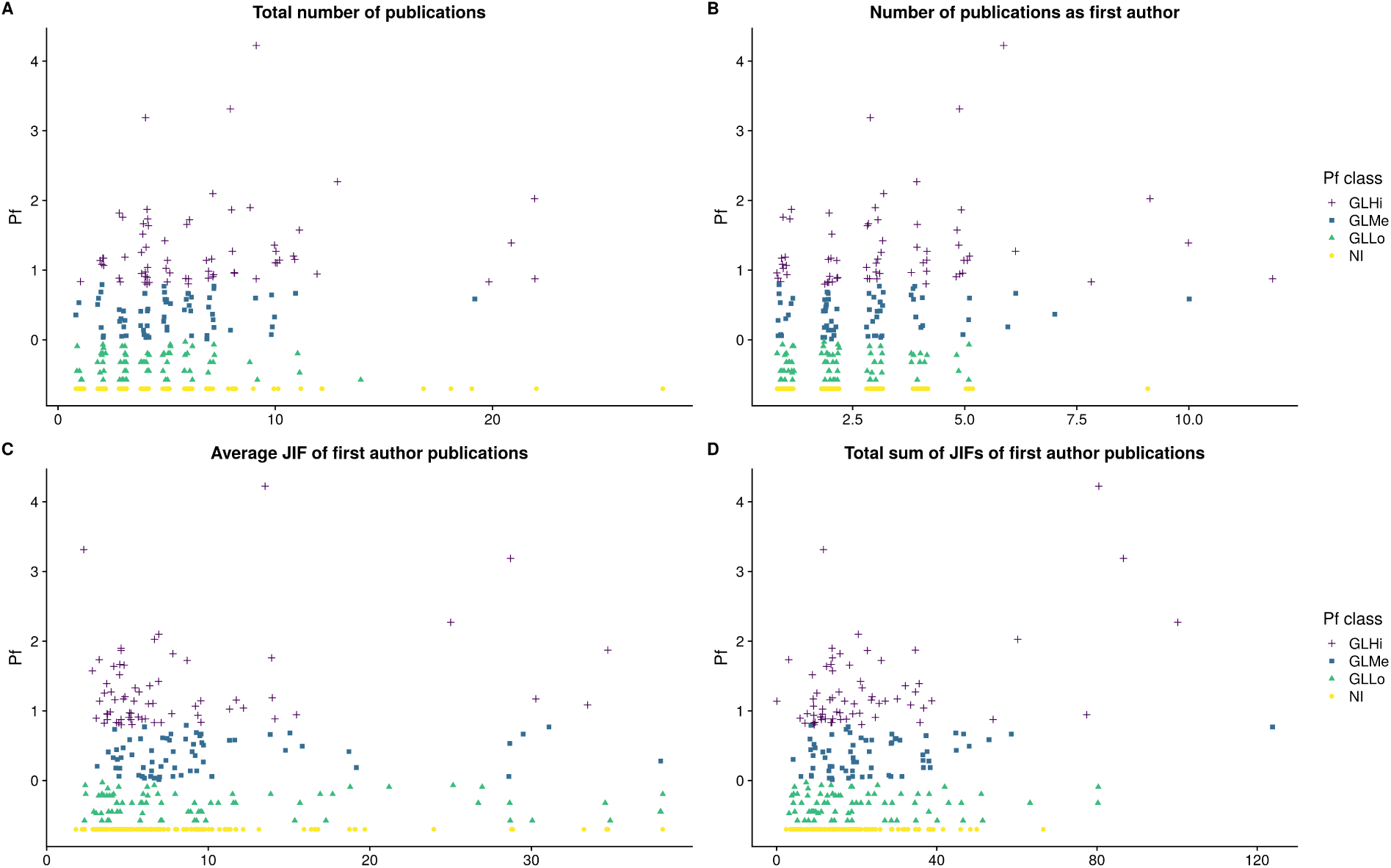
Relationships between scientific indicators and performance. (A) Performance factor and total number of publications. (B) Performance factor and first author publications. (C) Performance factor and average JIF of first author publications. (D) Performance factor and sum of the JIF of first author publications.

**Figure 3.**
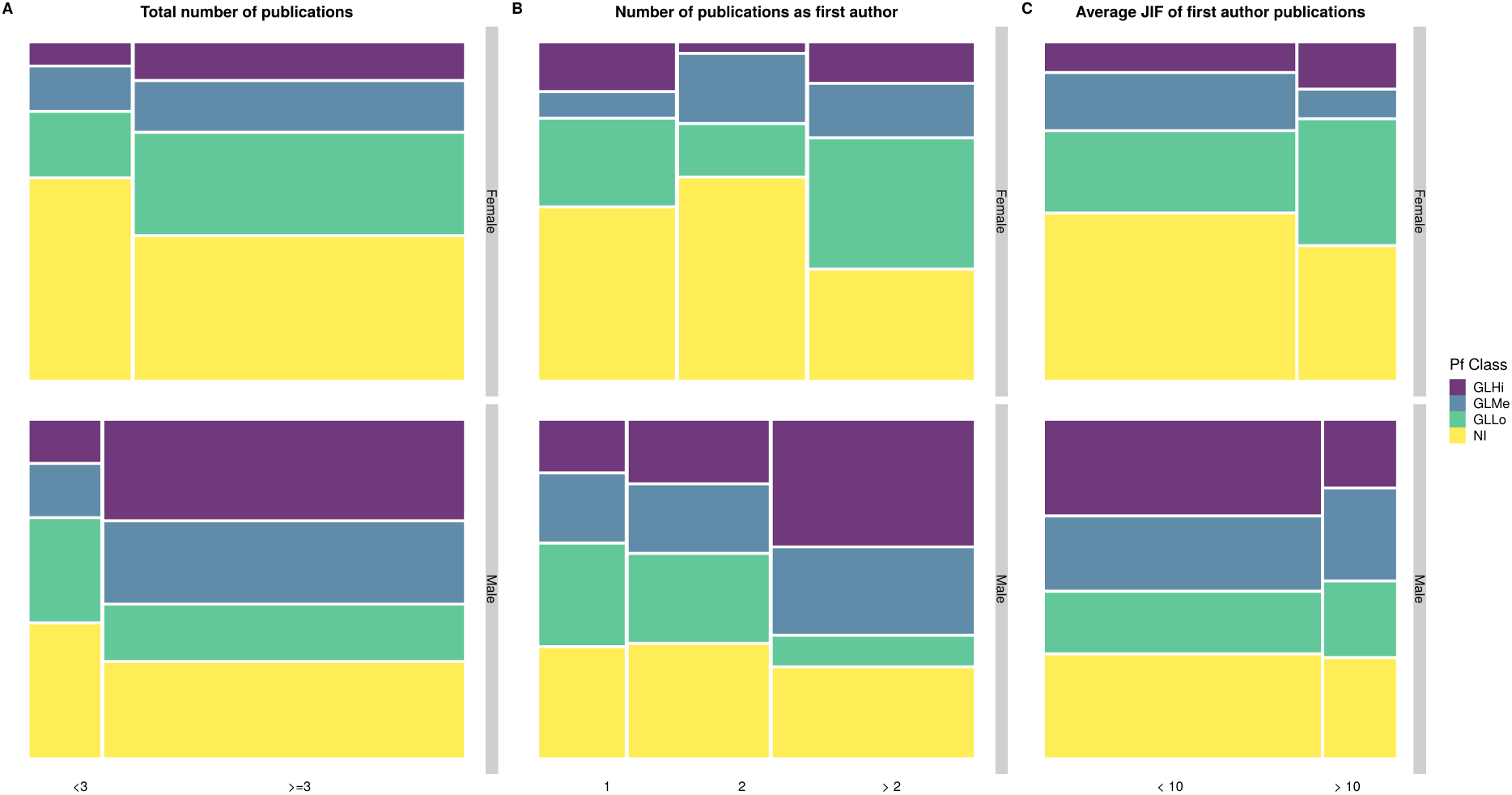
Mosaic plots for the relationships between scientific indicators and performace class. (A) Pf class and total number of publications. (B) Pf class and first author publications. (C) Pf class and average JIF of first author publications.

In particular, 3 or more total publications seemed to be associated with higher proportion of group leaders both in male and female candidates (Figure 3A), and this effect is clearer when only first author publications are considered, especially for female applicants (Figure 3B). However there were no statistically significant differences between male and female applicants in the number of first author publications in a log-linear model of homogeneous association controlling for Pf class (LR-Test, P=0.5668), as well as no association of a higher number of first author publications and a larger proportion of applicants who are group leaders in the same model (LR-Test, P=0.57555). Figure 3B indicates that there might be a small effect for female (and not for male) applicants. This small effect is however not statistically significant. In summary, a higher number of first author papers associates with a somewhat higher likelihood of becoming a group leader later on and this effect seems to be stronger for female than for male applicants, but this relatively small effect will require a larger sample to be confirmed or dismissed.

Dot plots were also generated for the distribution of candidates into Pf classes according to the average and sum of *JIFs* of the journals where first author articles were published (Figures 2C and 2D, respectively). Note that the pattern is less clear than for the number of publications and it is essentially random for the sum of the *JIFs*. Focusing on the average value of *JIFs*, values of average JIFs for first author publications were binned into two categories, above and below 10, and this was mosaic-plotted against gender and Pf classes. Figure 3C shows the average *JIF* is not significantly associated with the Pf class when controlling for gender (LR-Test, P=0.58732). Although there was a higher proportion of group leaders in the bin with an average JIF above 10 in female applicants, this difference was not statistically significant (LR-Test, P=0.22545).

In conclusion, analysis of the scientific indicators suggested that a higher number of first author articles published at the time of application mildly associates with a higher likelihood of becoming a group leader later on. However, this effect was essentially limited to female applicants and not statistically significant. Actually, it does not represent a large difference in absolute terms. For instance, the group of female candidates with only 1 first author publication is formed by 42 individuals, 22 of which are in the NI class. The group with more than 2 first author publications is formed by 51 female candidates, 17 of which are in the NI class.

Also interesting to note is the fact that despite the still pervasive misuse or JIFs as a proxy for individual performance and scientific quality in evaluation processes, we did not find any significant association between the JIFs of the first author publications published by candidates at the time of application and their future career progression according to Pf.

### The effect of social indicators on career progression

#### Supervisor networks and institutional prestige

As explained in the Methods section, the size of the scientific networks of the candidate’s PhD supervisor and the host supervisor (the principal investigator of the laboratory where the proposed project with be developed) were calculated using co-authorship as a proxy. The network size numbers used in figure 4 represent the actual number of unique co-authors in the 2 years prior to the first application deadline in 2007 (see **Methods**). Candidates were divided into two 5 groups according to network size, each group containing approximately the same number of individuals (see Figures 4A and 4B).

**Figure 4.**
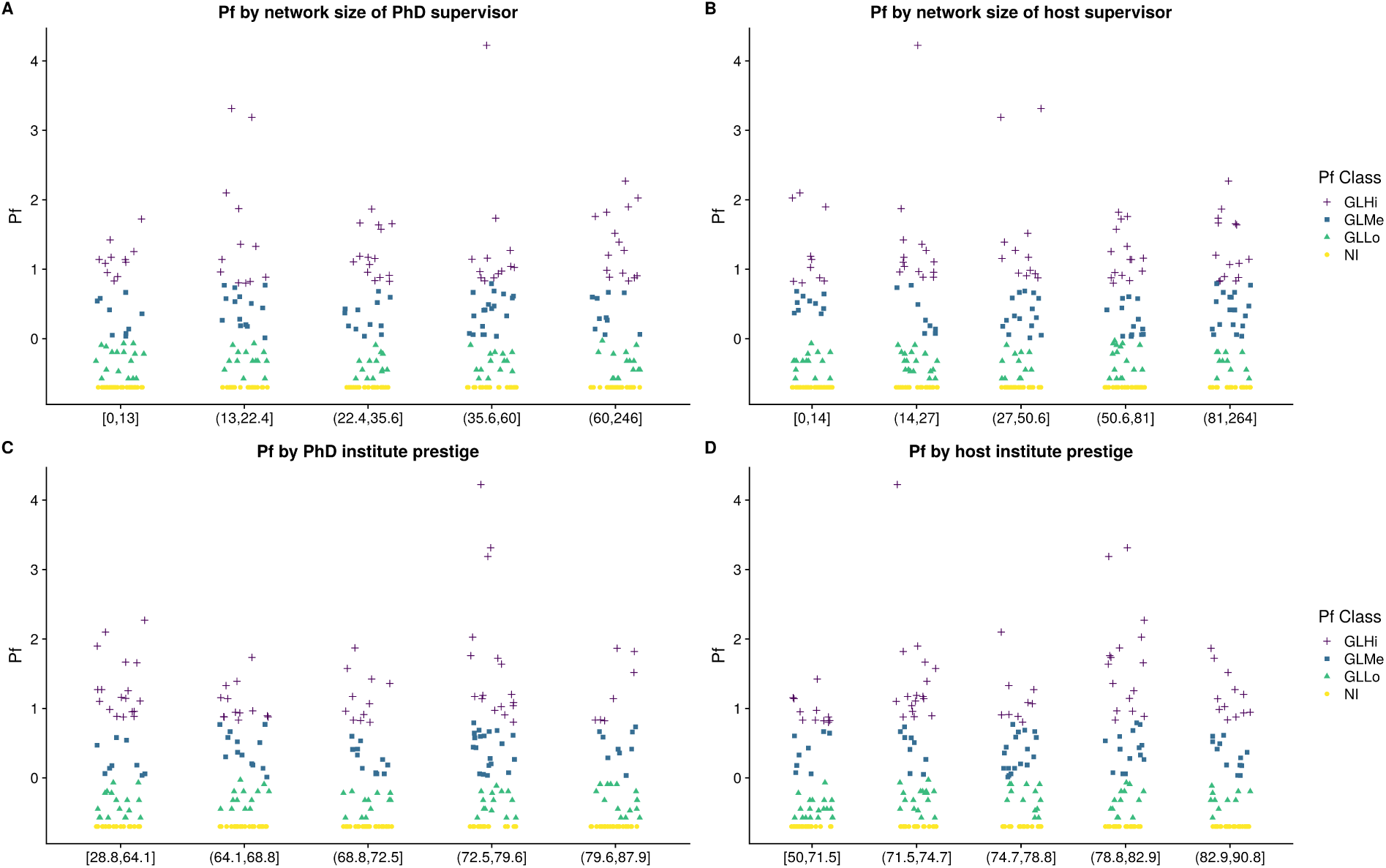
Performance stripcharts of social indicators. (A) PhD supervisor network size (B) Host supervisor network size. (C) PhD institute prestige. (D) Host institute prestige.

The analysis of the size of the networks of both the candidate’s PhD and host supervisor revealed rather similar distributions and did not show any statistically significant association with the distribution of candidates into Pf classes (ANOVA, P=0.69952 for the PhD supervisor network and P=0.84673 for the host supervisor network, excluding non-independent researchers for whom the Pf is constant).

Institutional prestige measured using a performance-based institutional ranking as a proxy (see **Methods**) did reveal a small effect (ANOVA, P=0.29439 for the PhD institution and P=0.032588 for the host supervisor institute). However, this was likely due to the effect of a small number of outstanding candidates who could be considered outliers. In fact, a Kruskal-Wallis test based on ranks did not reveal the same effect (P=0.52726 for the PhD institution and P=0.051632 for the host supervisor institute). For the analysis, candidates were again subdivided into 5 categories of roughly equal size based on the score *Best journal rate* provided by the Mapping Scientific Excellence application (see **Methods**). The higher the score, the higher the position in the ranking. No statistically significant association was found between institutional ranking and Pf classes. It is interesting to note, however, that most preselected candidates obtained their PhD in highly ranked institutions and plan to continue with their careers in also highly ranked host institutions, leaving less room for large differences. Also interesting is the fact that working in highly prestigious institutions does not seem to be per se predictive of future career progression: there was no general trend or strong correlation between prestige and Pf classes.

#### Nationality effect

Three different aspects were analyzed: applicant nationality, PhD country and host institution country.

Binning was again performed by dividing the group of applicants into 5 approximately equally sized groups (Figure 5).

**Figure 5.**
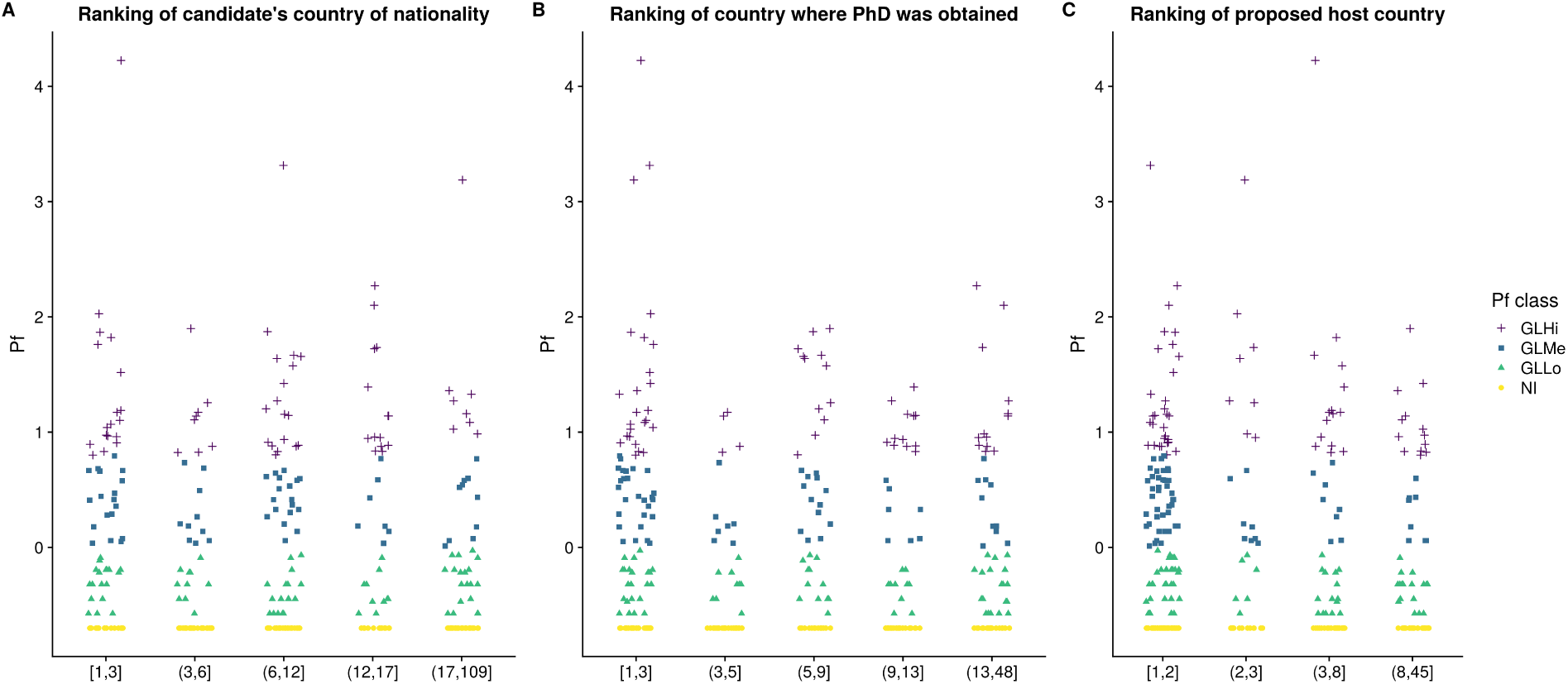
Performance stripcharts of nationality effects. (A) Candidate’s country of nationality (B) Country where the PhD was obtained. (C) Proposed host country.

Note that these semi-quantitative categories had different values for the different country rankings. Actual ranking positions were used in this analysis: the lower the value, the higher the position in the ranking.

Based on these binning criteria, rankings were analyzed with respect to Pf classes. Results suggest that country rankings (nationality of the applicant, country were the PhD was awarded and country where the host institution is located) are for the most part not associated to Pf classes. However, the country where the PhD was obtained and Pf classes seem to be somewhat associated, (Figure 5B; ANOVA, P=0.15652, again excluding non-independent researchers for whom the Pf is constant). However, Figure 5B suggests that this is mainly driven by a reduced number of outstanding candidates in the first bin combined with a below average performance in bin 2. Thus, this minimal, non-significant effect is probably highly specific to our dataset and may disappear in the noise of a much larger sample.

## Discussion

Peer review, the predominant mechanism of scientific evaluation at all levels, i.e. funding allocation, publication assessment or hiring decisions, functions adequately in gross discrimination of submissions, distinguishing between the flawed or inadequate and the high-quality ones. However, evidence provided in this study and elsewhere suggests that peer review fails to deliver precisely where it is needed the most nowadays: in finely discriminating among high-quality applications those that have a true potential from those that not (see **Introduction**).

We present here a detailed analysis of the peer review process among already preselected, high-quality applications to the ELTF programme with two questions in mind: are the decisions made by the peer review committee predictive of future scientific career progression? Are there indicators in the information presented at the time of application that may be predictive of future performance?

It is important to note that, besides scientific indicators traditionally used in scientific assessment, we have also analyzed what we refer to as “social indicators”. These are derived from information that is available at the time of application to the peer reviewers and that may be used consciously or unconsciously by peer reviewers in their decision making process: prestige (network size) of the PhD and host supervisors, country of origin of the applicant, country where the PhD was performed, country where the host institution is located, and institutions where the applicant has worked and plans to work to develop the proposed project (see **Methods** and **Results** for details).

Previous reports have certainly identified some objective positive correlations between these indicators and scientific performance. For example, scientific or personal communication among peers create social networks of scientists [34] that often result in collaboration in research projects and publication co-authorship [25, 26]. Network size can therefore be indirectly measured through co-authorship; increased co-authorship in turn, has been linked to increased productivity and impact [35–38]. Institutional rankings have also been found to be predictive of future performance and career progression [18, 27–29].

However, while this information may be part of the evaluation process, it is not always easily interpretable by evaluators and is a known source of bias. For instance, while most evaluators may concur on the importance of the number of publications during the PhD in the assessment, the relative importance of other factors such as the *prestige* of the host institution or the PhD supervisor is not clear and, in some cases may have unwanted effects on the decision.

Over-emphasizing the importance of supervisor networks or institutional prestige, may lead to well described decision biases such as the *Matthew effect*, according to which “eminent scientists get disproportionately great credit for their contributions to science while relatively unknown scientists tend to get disproportionately little credit for comparable contributions” [39; p. 56], or *affiliation bias*, which establishes that direct relationships between candidates (or candidates’ supervisors in this case) and evaluators has unwanted, positive or negative effects on the outcome of the evaluation [40, 41].

Similarly, while there is an expected effect of country scientific rankings on career progression based purely on differences in scientific productivity between countries [see for instance 42], there is also ample evidence of a direct effect of nationality, known as the *country-of-origin* effect, that can result in biases in decision processes. First described in the 60s in the field of Marketing, this effect is generally defined as the influence that knowing the country of origin of a product has on the consumer [43, 44]. Studies have shown that beyond quality considerations based on, to some extent objective data (analogous to scientific productivity, for instance), there are subjective values associated to countries of origin regarding economic status or exoticness, or linked to emotional connotations or even memories of past visits to those countries [for a review, see 45], all of which affect decisions.

### A failure of the peer review system?

In brief, the short answer to both questions above is no. No, ELTF awards are not predictive or future career progression and for the most part, no, the indicators tested are not predictive either. Maybe not surprisingly, this negative answer includes the predictive value of both social and scientific indicators, with the potential exception of the number of first author publications, which seem to have some small and non-significant predictive value of a better career outcome, particularly in female applicants.

Previous studies have described multiple reasons for evaluation failure resulting from the peer review process [7, 46]. As already mentioned, human bias is certainly one of the most studied and there are many factors, well beyond the scope of this article that, make human-based decisions markedly subjective. In the words of Cole and colleagues on applications to the National Science Foundation grants, “the fate of a particular application is roughly half determined by the characteristics of the proposal and the principal investigator, and about half by apparently random elements which might be characterized as ‘the luck of the reviewer draw’” [6: p. 885]. However, human bias is not the only potential source of error in the process. As already mentioned, evaluation criteria are not standard; there is no definition of the perfect scientific article, the perfect scientist or the perfect grant proposal; decisions are made by comparison among applications [4]. To make things worse, criteria for evaluation are not universal either, and even if they were, they are plagued with vague, difficult to objectivize terms such as “scientific quality”, “innovation” or “creativity” [47, 48]. Contrary to the scientific practice itself, scientific evaluation methods are based on notoriously vague concepts and therefore lack impartiality, understanding impartiality as “the ability for any observer to recognize evidence as evidence and to see the bearing of evidence on theory in the same way” [49: p. 352]. In line with this, a recent report on the peer-review process of NIH grant applications concluded: “results showed no agreement among reviewers regarding the quality of the applications […]. Although all reviewers received the same instructions on how to rate applications and format their written critiques, we also found no agreement in how reviewers “translated” a given number of strengths and weaknesses into a numeric rating” [50; p. 2952].

The vague definition of the criteria for evaluation only adds to the widespread use of single indicators in order to assess whether at least some of the evaluation criteria have been fulfilled [51], as already discussed above. Even nowadays, numerous funding agencies still require their applicants to list JIFs alongside publications and even if unasked, applicants often provide JIFs or equally simplistic factors such as h-index or number of citations as proxies for measures of productivity and scientific impact [24]. Our results suggest that these simple measurements of “scientific quality” have very limited predictive value of future performance once the top candidates have been preselected. In particular, despite the importance often attributed to JIFs, we did not find any association between the JIF of the journals where the candidates published the papers listed in their applications and their future scientific career progression.

An additional source of error sometimes overlooked in studies on peer review validity is intrinsic uncertainty. This refers to the fact that even if all biases could be eliminated, all the information available at the time of evaluation may be non-predictive of the future fate of the candidate or the project proposal and therefore may be insufficient to reach an informed decision. Our results for the ELTF selection process suggest that once a pre-selection has been done to exclude lower ranked candidates and a subset of highly competitive applications is under scrutiny, there is no evidence that the information available is sufficient to make valid decisions, at least when analyzed with the perspective of time. In fact, after the analysis of scientific and social indicators we can conclude quite the opposite: none of the indicators analyzed is predictive of future scientific career progression.

Obviously, we cannot exclude that other indicators not analyzed here would increase the predictive value of the information presented. Additionally, there are factors, such as the “quality”, “creativity” or “novelty” of the project presented by the applicant that cannot be directly measured but are used by the committee for the evaluation of proposals. That being said, the fact is that whatever information they used in their evaluation, the award of the fellowship itself is not predictive of future performance.

It should be pointed out that this study presents a rather detailed analysis of the application data and fate after 10 years of a group of applicants to the ELTF program. The level of detail attained, which is unusual in this type of studies, reduces substantially the number of individuals that can be scrutinized. Although the number analyzed here, 327 candidates (see the **Methods** section) is relatively large and allows for statistical analysis, it corresponds to a minority of the applicants to the program in a single year. Whether the results found here can be extrapolated to similar cohorts deserves further exploration in the future. Additionally, this study analyzes the relatively long-term professional fate of applicants, partly due to the fact that we wanted to explore career progression. We cannot exclude that analyses performed at different time points provide different results. For instance, indicators may be more predictive of performance at 5 years than they are at 10 years simply due to a reduction in the noise caused by external factors that affect scientific careers over time.

Other limitations are not particular to this study. For instance, the available tools have limitation themselves. There are no recognized standards for scientific evaluation. There are, as discussed, some measures that are reiteratively used in the literature and that we have explored in this study as well. However, the possibility exists that other indicators of, for instance, career progression or scientific quality, may render somewhat different results, although we believe that our conclusions at large would remain valid.

### Gender is predictive of career progression

Gender imbalance is a widespread issue in academia, and the difference between men and women becomes more obvious the more one individual advances in the academic career. For instance, in the EU-28 the proportion of STEM (Science, Technology, Engineering and Mathematics) female university students exceeds that of male students (59%) and the number of PhD graduates is rather similar within the two groups (47% of PhD students are women). However, further into the academic scientific career, numbers for women start to decline. Only 33% of researchers in the EU-28 countries are women, and only 21% of women occupy full professorships or equivalent positions (data from the European Commission (2015), *She figures 2015, gender in research and innovation*. Available at https://ec.europa.eu/research/swafs/pdf/pub_gender_equality/she_figures_2015-final.pdf, last accessed May 2018).

The situation in the United States is not very different according to studies performed in the last decade. As in Europe, about half of the doctorates in science and engineering are females, but women constitute only 21% of full science professors and 5% of full engineering professors [52].

The analysis presented in the present study further concurs with this data and multiple previous reports. The single most important predictor of career progression identified in this study is gender, and importantly, the gender gap observed according to Pf in 2017 is not explained by the relatively small differences in application and acceptance rates in 2007 (45% applications from women, 14% success rate for women, 18% for males). These differences are similar nowadays: aggregated application data from 2010 to 2016 reflects minor differences in the application and success rates between males and females (47% women applications, 13% success rate for women, 16% success rate for men, see http://www.embo.org/documents/news/facts_figures/EMBO_facts_figures_2016.pdf, last accessed May 2018). Gender has indeed been identified as a predictor of success in academic careers in other studies [see for instance 18] and numerous articles have reported gender inequality with regards to career progression in academia, including previous studies of the ELTF programme [53]. This effect has been shown to be stronger in STEM disciplines [54]. The reasons for this disparity are complex and well beyond the scope of this study. Particularly so when considering the fact that imbalance does not take place at the stage in which the object of this work, the ELTF programme itself, has influence on career progression (see above).

At the core of the causes of the gender gap observed among scientists, studies have identified gender bias, a male-centered working environment and naturally, work/family conflicts [55–58]. In a previous report on the ELTF programme performed in 2006, Ledin and collaborators surveyed candidates that applied in 1998. They found that although a comparable proportion of women and men had children in the period from 1998 to 2006 (61% and 69% respectively), only women reported having taken parental leave (2–3 months per child). Not only that, women were more willing to adjust their career for their families: they tended to move to suit their partner’s professional career more than men, they normally worked fewer hours per week than their partners and they usually contributed less than half of the income of the family [53].

Managing the work/family conflict is particularly important in academic settings, as the crucial period for career promotion is also the critical period for maternity in women. Career breaks or part-time work periods are very damaging at a time in life in which the publication record (productivity) is paramount for future opportunities [56, 58].

The period of 10 years used in this study from application to measurement of Pf fits well within the expected maternity window for women. Applicants were on average 30 years old at the time of application and more than 80% of the applicants in the analysis group did not have children at that time. However, demographic data on the applicants for the subsequent 10-year period is not available and further analysis is therefore not currently possible.

### Alternatives to peer review

In light of the issues discussed above, multiple alternative mechanisms have been proposed to allocate research grants in a more transparent and less biased manner. From a system in which all scientists are funded with equal amounts to systems based on pure scientific metrics to allocate funds, all have advantages and disadvantages [see for instance 14, 59, 60].

As already discussed, a generally accepted view is that a system that makes use of peer review selection will be able to gross discriminate among proposals, but fine selection is beyond its capabilities. Unfortunately, declining acceptance rates are forcing funders to make tough decisions precisely within the group where the peer review system has demonstrated its weaknesses, within the group in which differences between candidates cannot be identified and where decisions end up being opaque, biased and not anymore based on rational analysis.

Uncertainty and lack of objective factors to identify future high performers is not a problem exclusive of scientific selection processes. Human Resource Management literature also recognizes this problem in the selection of employees at large. Generally, if is admitted that future performance cannot be predicted with accuracy and, importantly, that lack of objective information for this prediction is often replaced by subjective opinions or biases [61, 62]. One well-documented example is what is called “statistical discrimination”: an employer might be prejudiced towards potential employees according to groups they belong to (black, women, Chinese, etc.) instead of focusing on their individual abilities for the job. For instance, previous good experiences with Chinese employees in a particular position, may create a bias, conscious or not, according to which being Chinese is a positive trait for that position [62, 63]. In order to minimize biases and noise in the selection process, industry has traditionally dealt with this uncertainty using a two-step process that is commonly practiced: first, selection processes are designed to extract as much information as possible from the candidates through tests and individual, panel and group interviews; and second, companies are allowed to real-life test their new employees during a probationary period to observe and evaluate actual performance [61]. Taking this strategy to the extreme, hiring practices such as “topgrading”, focused on the identification of the best possible candidates for every position, recommend a multistep hiring process, including several types of interviews, reference checks, analysis of the work and salary history of the candidates, and even coaching sessions [64].

Unfortunately, the economic costs, the effort and the time required to perform such in depth assessments makes these strategies unviable when more than five or ten candidates need to be evaluated. Fellowship or grant programs often face hundreds of applications. The study group presented in this report exceeds 300 candidates and the total number of applications for 2007 well exceeded 1000, well beyond the capabilities of any organization to use strategies such as topgrading.

In essence, uncertainty, lack of objective indicators and bias preclude a system based on peer review to faithfully rank these 300 candidates. Once the weaker applications have been excluded in a pre-selection step, the process of selection of awardees is statistically indistinguishable from random allocation of awards if the validity of the selection is evaluated with the perspective of time as we do here. Except that allocation is not random, it has a significant component of uncertainty, lack of information and more importantly, biases. “Scientists are human, and thus susceptible to biases. […] When there is no objective basis for choosing on candidate over others, people naturally fall back on subjective preferences. A selection committee might consciously or unconsciously favor certain research topics, groups of people or even individuals” [15; p. 7].

### Living with uncertainty: focal randomization

If we admit that peer review evaluation fails at distinguishing applicants with a high potential from those with slightly less potential or even worse, that current success rates are causing the exclusion of candidates just as good as the ones selected, we have to conclude that once pre-selection has taken place, final allocation of grants is not based on objective, predictive criteria. On the contrary, selection is more similar to a random process, but potentially plagued with biases or personal preferences. However, if selection resembles a random process, why not making it truly random to avoid any and all potential biases?

We believe that a system that combines the peer review system with a randomization step to allocate funds among those applications that cannot be further distinguished according to the criteria used in their evaluation, represents a honest and transparent selection procedure based on the evidence available.

Top-ranked applications would be funded, applications deemed of low quality would be denied funding and applications for which this discrimination cannot be made would enter a process of random allocation of funds. Randomization would therefore be *focal*, as it would only be applied to a number of applications selected by the peer review committee, which would be otherwise forced to make decisions on applications in which peer review has consistently shown weaknesses [65].

It is important to emphasize that this procedure would not replace peer review with random allocation of funds. It would leave peer review at the position where rational decision has been shown to perform at its best, and use focal randomization to allocate grants beyond the limits of peer review.

---

**Supp. Figure 1.**
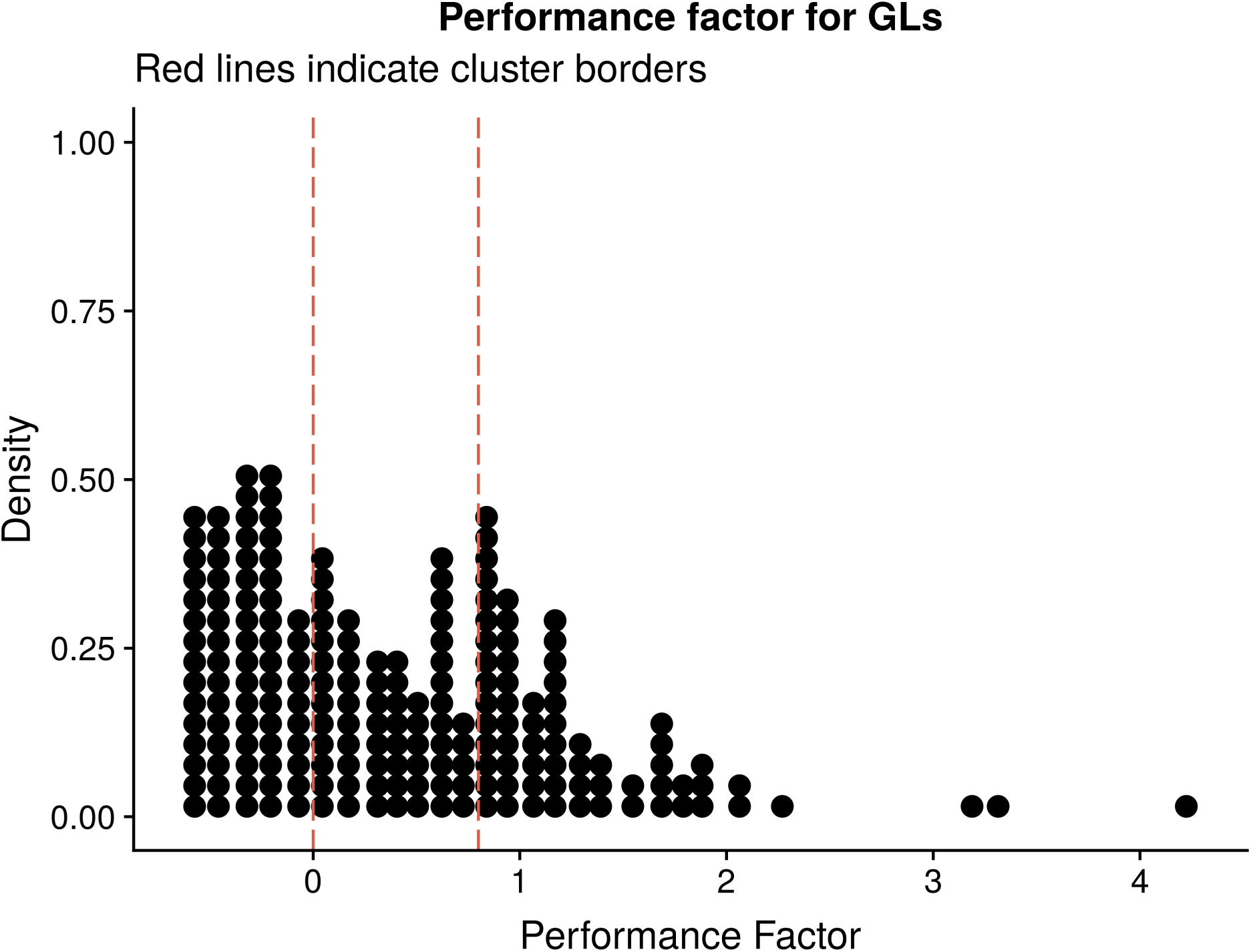
Dotplot representation of the Pf binning.

**Supp. Figure 2.**
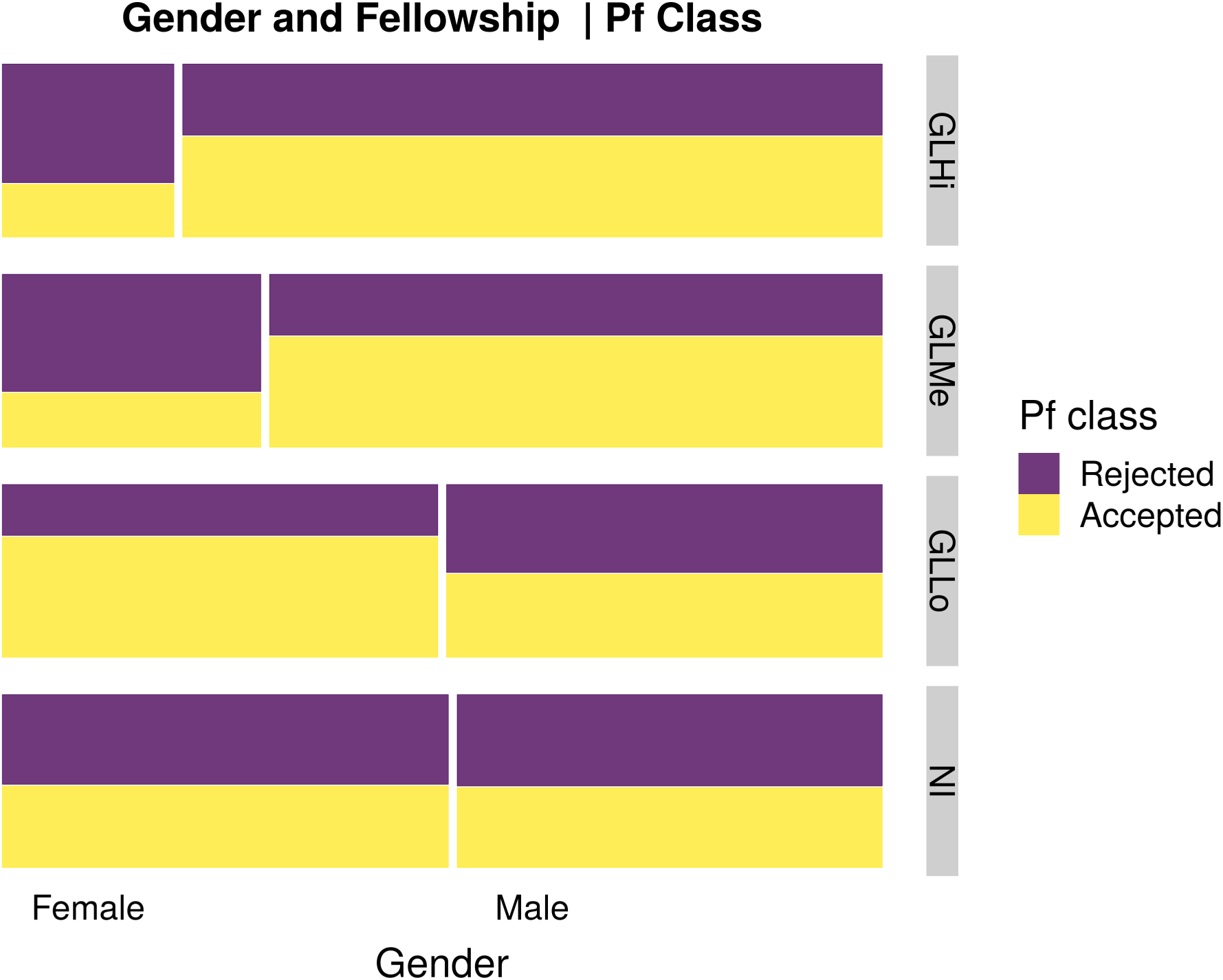
Mosaic plot of the relationship between gender and acceptance into the fellowship given the Pf class.

